# Interplay of pupil-linked arousal and cortical decision computations in mice

**DOI:** 10.64898/2026.06.05.730199

**Authors:** Maxime Maheu, Jasper B. Copeland, Alexis Cerván, Jaime de la Rocha, J. Simon Wiegert, Tobias H. Donner

## Abstract

Influential frameworks hold that brainstem neuromodulatory systems are rapidly recruited in a top-down manner during decision-making, adapting brain-wide activity and cognitive behavior in real time. Here, we examined the relationship between decision-related frontal cortical activity and pupil responses (a peripheral proxy of neuromodulatory activity) in mice accumulating fluctuating auditory evidence to choose among two options. We found two components of preparatory activity during decision formation, prior to motor action: a non-selective component shared between hemispheres and a choice-selective component that reflected decision uncertainty. Both components were mirrored in pupil responses during the task, which, in turn, scaled with decision uncertainty during decision formation and with uncertainty-modulated prediction errors after experiencing the outcome. Pupil responses also predicted an adjustment of choice behavior at the next trial. Our results point to a recurrent interplay between cortical decision computation and pupil-linked neuromodulation.

**HIGHLIGHTS:**

- Mice switch between evidence integration and heuristic strategies during auditory-informed decisions
- Pupil responses track decision uncertainty and uncertainty-modulated prediction errors
- Decision-related frontal cortical activity is coupled with pupil responses
- Pupil responses predict an adjustment of future choice behavior

**IN SHORT:** Maheu et al. show that the phasic transitions in the arousal state mirror distinct components of cortical activity, including one directly involved in decision computations. In turn, these pupil-linked arousal responses predicted the strategy adopted at the next trial.

## INTRODUCTION

Adapting to environmental contingencies requires the coordination of computations across brain-wide networks. A seminal hypothesis posits that key cognitive quantities computed in higher-tier brain regions involved in inference and decision-making are broadcast across the forebrain by an evolutionarily ancient arousal brainstem system.^1–3^ This system comprises chemically distinct neuronal populations with widespread axonal projections through which they release modulatory neurotransmitters. The latter are capable of altering cortical computation by shaping synaptic transmission^4^ and plasticity.^3,5^ Conversely, many of these brainstem arousal centers, including the noradrenergic locus coeruleus, receive top-down projections from the cortical regions they modulate.^1,6^ This architecture establishes a large-scale loop in the mammalian brain enabling local cortical decision computations to shape brain-wide activity and cognitive behavior.

It has been proposed that during cognitive processing, neuromodulatory systems are recruited by latent variables reflecting ongoing decision computations via frontal cortical regions.^1^ Specifically, the anterior cingulate cortex may compute a conflict signal that reflects the strength of competition between neural populations encoding different choices (neural ‘decision variables’) during decision formation.^7^ At the same time, the orbitofrontal cortex may use these neural decision variables to compute decision confidence, or its complement, uncertainty.^8,9^ Such uncertainty signals, in turn, translate into graded prediction errors during experience of the decision outcome.^10^ Together, these latent computational variables may shape rapid variations in arousal state during task performance. However, direct empirical support for this integrated framework remains lacking.

Non-luminance-mediated fluctuations in pupil size are frequently used as a peripheral proxy of central arousal state.^11^ Indeed, pupil dynamics closely track the activity of the locus coeruleus^6,12^ and dorsal raphe,^13,14^ alongside other neuromodulatory nuclei such as the basal forebrain,^15^ nucleus incertus^16^ and hypothalamus.^17^ Accordingly, pupil responses in humans also reflect uncertainty during decision formation and uncertainty-modulated prediction errors after outcome delivery.^18,19^ The relationship between specific cognitive variables and pupil-linked arousal has been studied primarily in humans and only rarely in mice.^20–22^ This represents a critical gap given that the mouse brain serves as the primary model system for dissecting neuromodulatory circuits^12–15^ and brain-wide cognitive computation.^23–25^ Specifically, while recent work has linked baseline pupil sizes before task execution to overall task engagement in mice,^22,26^ it remains unknown whether and how task-related pupil responses track specific computational variables such as decision uncertainty. Most fundamentally, it has remained unclear whether and how these phasic pupil responses during decision-making relate to decision-related cortical signals. The present study aimed to close these gaps.

We monitored pupil responses in mice performing a challenging two-alternative forced choice auditory decision task that promoted the protracted accumulation of fluctuating sensory evidence. Concurrently, we measured the activity of bilateral premotor cortex (anterior lateral motor cortex, ALM) during decision formation.^27^ We observed that phasic pupil responses scaled with decision uncertainty. Critically, moment-to-moment variations in two distinct frontal cortical signal components – a choice-selective and a non-selective component – correlated differently with these pupil responses. The pupil dynamics, in turn, predicted adjustments in future choice behavior. Our results demonstrate a tight interplay between cortical decision computations and pupil-linked arousal systems in the support of adaptive cognitive behavior.

## RESULTS

### A behavioral task dissociating decision formation from motor report and feedback

We trained head-fixed mice in a novel behavioral task which promoted the integration of fluctuating auditory evidence over time (**Fig. 1a**). Specifically, mice listened to a 1s-long, binaural sound delivered from speakers on the left and right sides before they were prompted to report the side associated with the loudest speaker on average. They did so by licking one of two ports mounted on a linear actuator, which kept lick ports beyond reach for a delay period of 1s from evidence offset (see **Methods**). This delay decoupled the processing of the sensory evidence and concomitant decision formation on the one hand, and the execution of the motor act used for reporting the choice (lick) and the processing of the choice outcome (reward or no reward).

**Figure 1.**
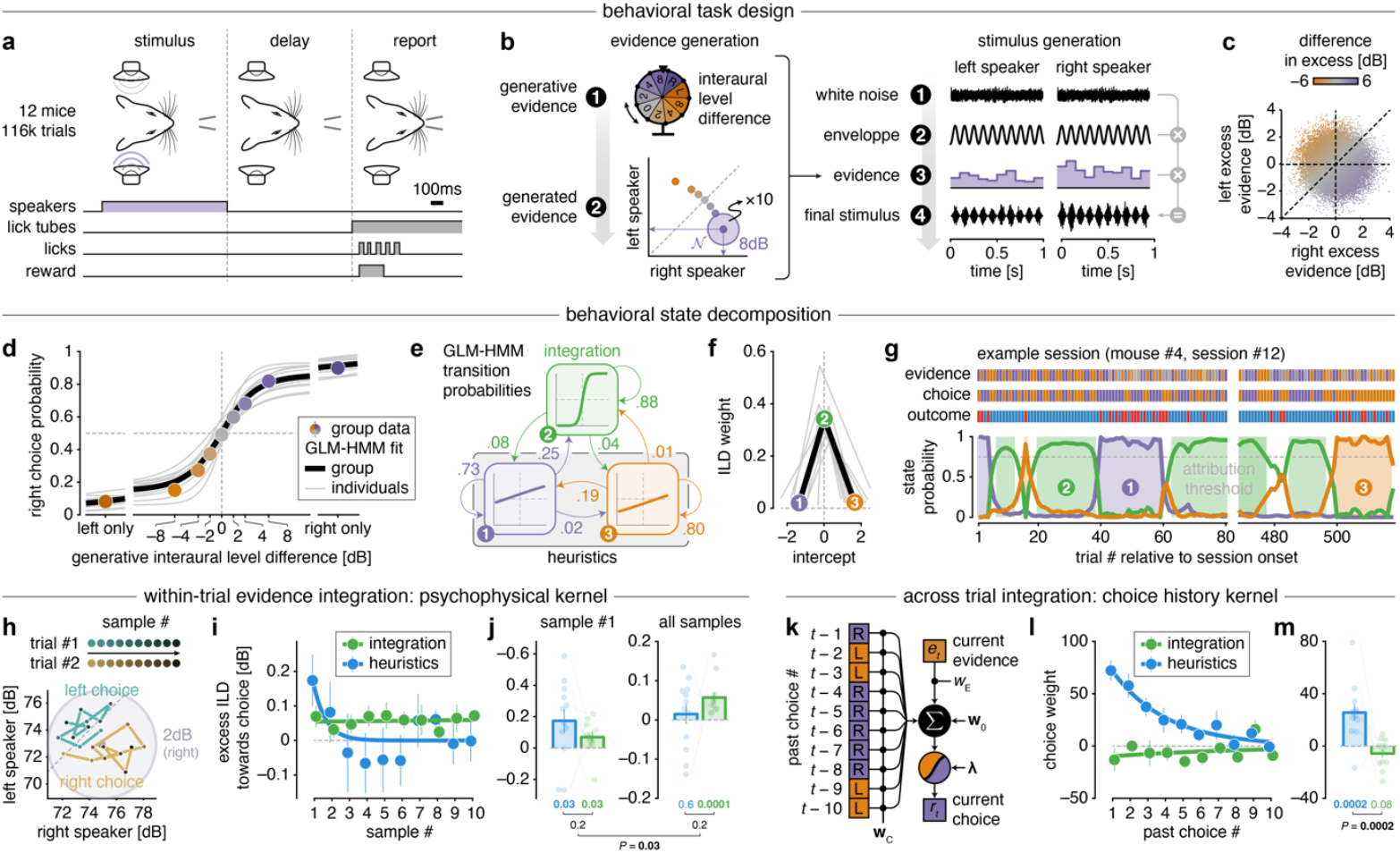
Mice integrate auditory evidence in a behavioral task dissociating decision formation from motor report. **a**, Head-fixed mice are trained to collect a water drop on the side (left or right) associated with the loudest speaker. Lick tubes are only accessible after the stimulus is played (1s) and a delay period has elapsed (1s). **b**, At each trial, the random draw of a generative evidence level determines the relative intensity of left and right speakers (generative interaural level difference = 0, ±2, ±4, ±8dB, or single-speaker denoted by L and R). Positive ILD indicates that the right speaker is louder than the left, and negative ILD to the converse. Note that generative probabilities were not balanced and easy trials were more frequently sampled to maintain engagement. Ten individual samples are drawn from a normal distribution centered on the chosen generative ILD. The stimulus consists in white noise bursts whose amplitude is scaled by each of the 10 stochastic evidence samples. **c**, Individual samples used in the experiment after subtraction of mean ILD (corresponding to ‘excess evidence’ in panels h-i). **d**, Group mean ± s.e.m. right choice probability as a function of ILD and fit by a generalized linear model (GLM) hidden Markov model (HMM) with 3 states. Thin lines indicate fit to individual mice. **e**, Schematics of the GLM-HMM along with estimated transition probabilities between each pair of states. **f**, Group mean ± s.e.m. GLM parameters in each state (sorted). **g**, Snippet of generative ILD (same color code as in panel b), choice (purple: right, orange: left), outcome (blue: correct, red: error) and corresponding state probability from an example session in an example mouse. A state probability threshold of 0.75 is used to allocate trials to integration-(green) or heuristic-based states. **h**, Two example trials generated from the same generative ILD (2dB) leads to different choice because of stochastic fluctuations. This feature allows to infer the respective use of each of the 10 samples (using excess evidence in panel c). This analysis was restricted to 0-4dB generative ILD as these are the only conditions with incongruent samples relative to the generative mean. **i**, Group mean ± s.e.m. of psychophysical kernels, defined as excess evidence in chosen (positive) vs. unchosen (negative) options for each sample position, sorted by behavioral state. **j**, Group mean ± s.e.m. difference in excess evidence at first sample (left) and averaged across all samples (right). Dots and thin lines correspond to individual mice. *P*-values are obtained from a random permutation test (10,000 permutations). **k**, Schematic of the choice history logistic model. **l**, Group mean ± s.e.m. regression coefficients (from model in panel j) for past choices, separately for each behavioral state from the GLM-HMM (left). **m**, Same as panel i but for average choice weight in panel k (right).

Evidence strength was defined by the inter-aural level difference (ILD; (**Fig. 1b-c**), which we varied from trial-to-trial: from ‘ambiguous’ (ILD = 0dB) to ‘difficult’ (±2, ±4dB), ‘easy’ (±8dB), and ‘trivial’ in which only one speaker was active (ILD = ±80dB). Each sound was composed of 10 independent white noise bursts (‘samples’) whose intensity (i.e., loudness) was stochastically drawn from a distribution centered on the generative ILD. Performance on trivial trials (ILD = ±80dB) was not limited by perceptual uncertainty and thus used to assess task comprehension and engagement. In these trials, mice reported 91% (range: 85-96%) decision accuracy. On trials where both speakers were active, choice probability scaled lawfully with the signed evidence strength in all mice (**Fig. 1d**), indicating that their behavior was, on average, under stimulus control.

### Mice alternate between integration and heuristic-based strategies

The characterization outlined above captured the animals’ task performance aggregated across the entire session. Recent modeling work has shown that mice tend to switch between different (task-engaged versus disengaged) states throughout the protracted performance of a perceptual choice task.^28^ To uncover such putative state-switching, we fitted a logistic choice model whose parameters were free to vary between discrete states according to a 3-state hidden Markov process (**Methods**). This approach identified one stimulus-dependent (i.e., ILD weight >> 0) state and two other states dominated by streaks of either left or right choices (i.e., |intercept| >> 0; **Fig. 1e, f**). These locally biased states were evident by inspecting individual sessions (**Fig. 1g**). Mice tended to remain in the current state and occasionally transitioned to another of the two states. We grouped biased states together and used a 0.75 state probability threshold to label trials according to state identity,^29,30^ yielding a dataset comprising 55% of trials with a response (range: 20-83%). We deliberately avoid labelling these states in terms of task engagement, as the mice remained highly engaged throughout the session, reporting decisions in 99% of trials (range: 98-100%).

The states were associated with distinct decision-making strategies, which we refer to as ‘evidence integration’ and ‘heuristics’ in the following (**Fig. 1h-m**). To estimate individual evidence integration profiles, we leveraged trial-to-trial fluctuations in the evidence on trials with ambiguous or very weak evidence (0-4dB). We computed so-called psychophysical kernels^31–33^ in terms of the excess evidence (difference in sample-wise ILD from mean ILD) toward the chosen option (**Methods**; **Fig. 1h-i**). Excess evidence greater than zero at a given sample position indicates that evidence fluctuations at that samples affected choices. In the biased states, only the first sample had a strong contribution to choice, with close to zero excess evidence for all remaining samples (**Fig. 1h-i**), indicating the absence of integration of all available evidence. In contrast, in the stimulus-dependent state, the excess evidence was greater than zero for all samples. These flat psychophysical kernels are in line with a protracted integration of all evidence throughout the trial.

The two states were further dissociated in terms of the relative contribution of history of past decisions on the current choice (**Fig. 1k,l**). A tendency to repeat previous perceptual choice, even when (as in our task) stimulus sequences are random across trials, is pervasive in rodents^34–36^ and humans.^37^ We used logistic regression to quantify the biasing effect of previous choices on current choice (**Fig. 1k**; **Methods**). As expected, the biased state was characterized by substantial effects of previous choices which gradually decayed as a function of lag from the current trial (**Fig. 1l**). Indeed, this is another reflection of the same local side bias identified in **Fig. 1f**. Critically, however, we found that such choice history bias was nearly absent from mice behavior in the stimulus-dependent state.

Taken together, the analysis of psychophysical kernels and choice history effects indicate mice employed a strategy that is maximally in line with task demands in the stimulus-dependent state. These findings contribute to the growing realization that mice exhibit near-optimal accumulation of evidence for decision-making under uncertainty.^24,38^

### Behavioral and pupil signatures of decision uncertainty

Decision-making under perceptual uncertainty is associated with a level of confidence, the probability that a choice is correct given the available evidence.^9^ In tasks where the evidence duration is fixed, such as the one used here, a hallmark signature of decision confidence (and its complement, uncertainty) is an opposite-signed scaling with evidence strength on correct versus error trials.^8^ This signature can be explained with a simple model, in which confidence is determined by the absolute distance of a decision variable at the time of choice (**Fig. 2a**). Across rodents, non-human primates, and humans, neural activity and behavior – including reaction time (RT) – before feedback exhibits this signature in many contexts.^8,10,18,19,39^

**Figure 2.**
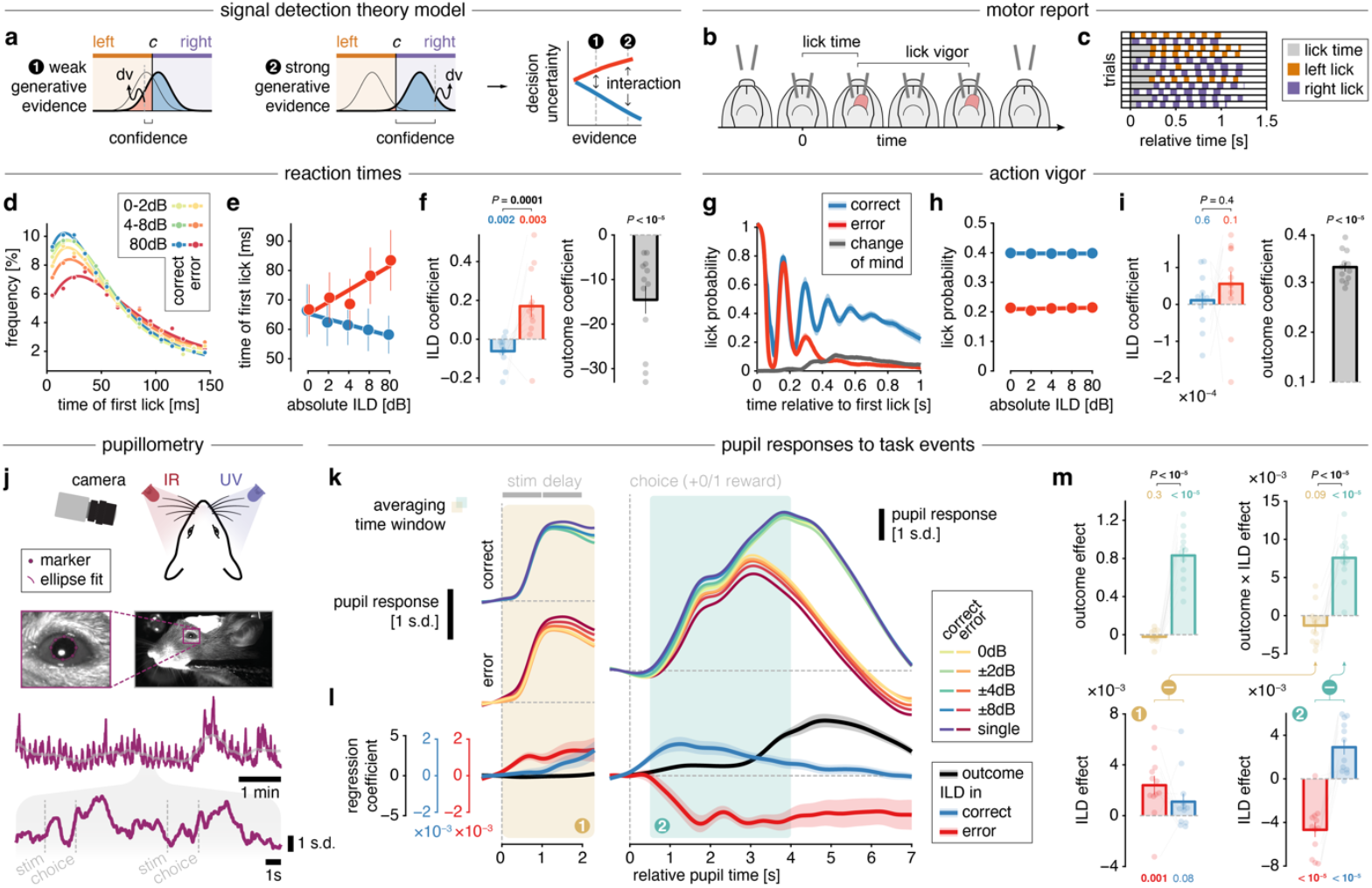
Behavioral and pupil signatures of decision uncertainty. **a**, Schematics of a signal detection theory model of decision uncertainty (left: 1, weak evidence; right: 2, strong evidence). The internal decision variable (dv) on each trial is a sample drawn from the corresponding distribution (shown here for a Gaussian). It is compared with a decision criterion *c*, to compute a binary choice as well as a graded measure of decision confidence. Decision uncertainty (1-confidence) scales with evidence strength with a sign that differs by outcome. **b**, In the task, reaction time corresponds to the time between when lick tubes become available and the first lick while lick vigor corresponds to lick frequency during a 1s time-window following the first lick. **c**. Example trials depicting lick micro-structure. **d**, Distribution of reaction times as a function of binned inter-aural level difference (ILD) and outcome. Dots correspond to data in 15 equally-spaced bins and lines to a log-gaussian fit. **e**, Group mean ± s.e.m. of the median time of first lick according to absolute inter-aural level difference (ILD; x-axis) and outcome (color coding). **f**. Group mean ± s.e.m. regression coefficient of ILD separately in each (left) and for outcome (right). *P*-values are obtained from a random permutation test (10,000 permutations). **g**, Group mean ± s.e.m. lick probability aligned to the initial lick. Traces are categorized by whether the first lick was on the correct or error side. The ‘change of mind’ trace represents licks on the tube opposite to the one chosen initially. **h**, Same as panel e but showing the average lick probability in a 0-1s time-window following the first lick. **i**, Same as panel f but for lick probability shown in panel h. **j**, Eye is video monitored using an infrared (IR) camera and a contralateral ultraviolet (UV) light source maintains a stable luminance environment while maximizing dynamic range of pupil fluctuations. Pupil size corresponds to the diameter of an ellipse fitted on the pupil edge markers (top). Snippet of pupil size and zoom-in in two example trials (bottom). **k**, Group mean pupil response to stimulus (left, split by correct vs. error) and choice (right). Shaded areas correspond to time-windows for averaging and group-level statistics. **l**, Group mean ± s.e.m. time-course of correlation coefficient quantifying the effect of outcome and inter-aural level difference (ILD; separately on correct vs. error trials) on pupil responses. **m**, Group mean ± s.e.m. effect of outcome (top left), ILD (bottom), and their interaction (top right). *P*-values are obtained from a random permutation test (10,000 permutations).

Consistently, mice in our task exhibited this identical behavioral pattern (**Fig. 2b-f**). Given the imposed delay, the animal’s choice reports were typically triggered by the presentation of the lick tubes (**Fig. 2b**). We defined RT as the latency of the first lick (**Fig. 2b-c**). Distribution of these latencies exhibited the skew commonly observed for RT measures (**Fig. 2d**). Analyzing mean RTs as function of evidence strength and outcome revealed the interaction expected for decision uncertainty (**Fig. 2e-f**), precisely as predicted by the uncertainty model (compare with **Fig. 2a**). In these analyses, we used all trials irrespective of strategy to ensure sufficient trial counts for errors on easy trials (**Methods**).

While mice were uncertain about the correct choice at the time of their first lick, water delivery (or absence thereof) was determined by that lick (**Methods**). Thus, subsequent licking behavior should be informed by the outcome. Lick vigor exhibited a commonly observed 7 Hz oscillation^40^ (**Fig. 2b-c,g**). While it was stronger for correct than incorrect responses, it was independent of evidence strength (**Fig. 2h-i**). In other words, lick vigor reflected the complete resolution of decision uncertainty provided by the outcome.

Having established that animal’s behavior (RT) was shaped by decision uncertainty, we next tested whether the same held for a measure of arousal: the non-luminance mediated changes in pupil size. We video-monitored the size of the left pupil while mice were engaged in the task and analyzed pupil responses during decision formation as the baseline-corrected change in pupil size evoked by the onset of the auditory stimulus (**Fig. 2j**; **Methods**). The delay separating evidence and choice report allowed us to dissociate the scaling of pupil responses before uncertainty resolution and compare it to responses following its resolution, then time-locked to the first lick.

The pupil dilated following evidence onset (during decision formation) and, more strongly, following the choice report (associated with outcome delivery and ongoing licking behavior; **Fig. 2k**). As for RTs and lick vigor, we analyzed pupil responses in these two time windows as a function of evidence strength and outcome (correct versus error).^5,6,29^ For each time window, we observed the interaction between the two factors predicted by the uncertainty model, but with a sign flip between the two time windows (**Fig. 2l**). Such a sign flip of the interaction between evidence and outcome has been previously observed in the activity of dopaminergic midbrain neurons in monkeys^10^ and in pupil responses in humans.^18^ Different from uncertainty model predictions (**Fig. 2a**) and RTs (**Fig. 2e**), the pupil responses scaled positively with evidence during decision formation (first time window) for both error and correct trials. The positive slope before correct choices may reflect a positive reward expectation component superimposed on a negative uncertainty component. Critically, as predicted by the model, the slope of the scaling was larger for error trials (**Fig. 2l,m**). The flipped pattern of the slopes for correct and error trials after outcome delivery (second time window, **Fig. 2l,m**) is consistent with an uncertainty-modulated reward prediction error signal that has so far only been observed in primates.^10,18^ There was a main effect of outcome on pupil responses only in the second time window (**Fig. 2m**), which may reflect the difference in licking behavior between correct and error trials (**Fig. 2**).

In sum, pupil responses during different stages of the task showed an intricate pattern of evidence- and outcome-dependent modulations that reflect uncertainty-modulated internal decision variables (before choice) as well as uncertainty-modulated prediction errors (after choice) known to exist in the primate brain. Together with our characterization of choice behavior (**Fig. 1**), this indicates that mice’s decision behavior and the concomitant recruitment of their arousal system, followed principles of normative probabilistic inference. We next tested for cortical signatures of decision formation, which may be involved in the recruitment of pupil-linked arousal.

### Two components of premotor cortical activity during decision formation

We recorded the cortical population signatures of decision formation in a subregion of the antero-lateral motor (ALM) cortex. The ALM is analogous to the premotor cortex in primates exhibiting ramping signals that track latent decision variables during evidence accumulation^,27,41,42^ and which hosts choice-selective neurons.^43^ While cells in the ALM that prefer left- and right-choice are intermingled in both hemispheres, they exhibit a biased representation at the level of the population such that each neuronal group would predominate in the hemisphere contralateral to choice. This should enable choice-selective activity to be tracked in mesoscopic population signals from left and right ALM, analogous to non-invasive approaches in humans.^27,32,41^

Given ALM sends callosal fibers to the contralateral hemisphere^,44^ we monitored the left and right ALM with spectrally distinct genetically encoded calcium indicators (jGCaMP8m and jRGECO1a; **Fig. 3a**) to avoid cross-talk between hemispheres. We focused on ALM population dynamics during decision formation following evidence onset (stimulus and delay intervals), but before the licking interval that was dominated by vigorous motor movement. On average across all trials, ALM activity exhibited similar dynamics in both hemispheres, which was characterized by an initial increase upon evidence onset, followed by a prolonged increase throughout the rest of the stimulus and delay intervals (**Fig. 3b**).

**Figure 3.**
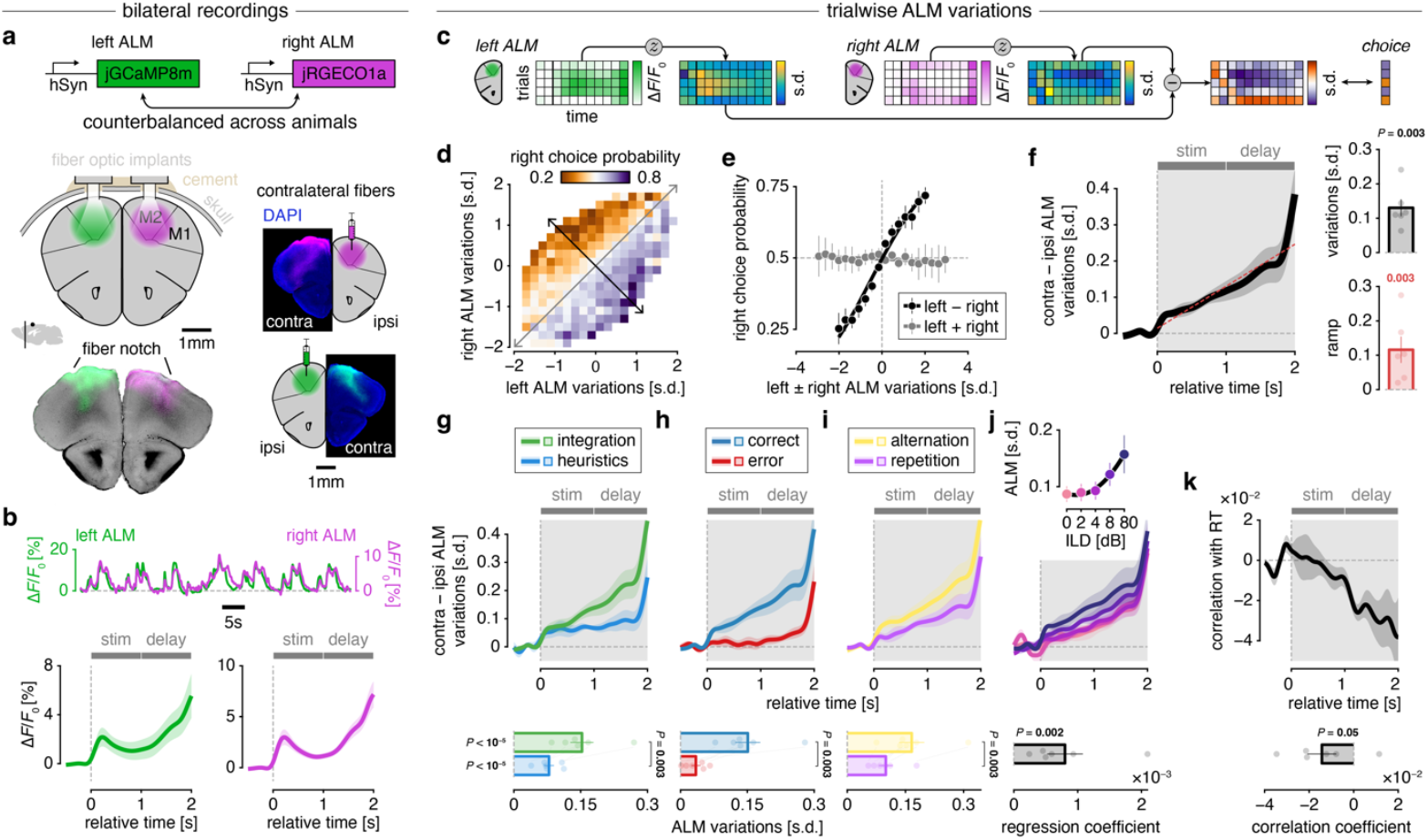
Bilateral population activity in premotor cortex entails decision-independent and decision-related signals. **a**, Expression of jGCaMP8m and jRGECO1a in left and right antero-lateral motor (ALM) cortex is counter-balanced across mice. The use of spectrally different calcium indicators is motivated by the existence of callosal fibers to the contralateral hemisphere and allow to record pure unilateral signals. **b**, Snippet of left and right ALM activity in an example mouse and corresponding (top). Group mean ± s.e.m. left and right ALM response to stimulus (bottom). **c**, Trial-by-trial fluctuations of single-hemisphere activity is standardized across trials (at the session level) and standardized activity in hemisphere ipsilateral to subsequent choice is subtracted to that of the contralateral hemisphere. **d**, Group mean right choice probability as a function of left and right ALM standardized activity averaged during stimulus and delay periods. **e**, Group mean ± s.e.m. right choice probability as a function of the difference or sum of left and right ALM standardized activity. Lines correspond to sigmoid fits. **f**, Group mean ± s.e.m. difference in standardized ALM activity contralateral vs. ipsilateral choice aligned on stimulus onset (left). Barplots show the average in stimulus and delay periods (top) and the slope of the ramp measured in the same time window (bottom). *P*-values are obtained from an exact permutation test. **g-j**, Group mean ± s.e.m. contralateral vs. ipsilateral ALM variations (as in panel f) but split according to different trial types. Barplots show the average in stimulus and delay periods and *P*-values are obtained from an exact permutation test. **k**, Group mean ± s.e.m. correlation coefficient between contralateral vs. ipsilateral ALM variations at each time-point and reaction times (time of first lick).

To isolate choice-selective activity, we computed the difference between standardized ALM activity contralateral and ipsilateral to the upcoming choice (**Fig. 3c**; **Methods**). The resulting signals co-varied across hemispheres and contained two components related to the task: their hemispheric difference (i.e., lateralization) which predicted choice and their sum (i.e., ‘global’ component), which did not entail choice information (**Fig. 3d,e**). Similar orthogonal components of task-related activity have been observed in primate motor and visual cortex.^27,45–49^ The choice-selective ALM component also gradually (and approximately linearly) built up during evidence integration followed by a further transient right before choice report (**Fig. 3f**).

The choice-selective component of ALM dynamics mirrored several key aspects of behavior. First, it only ramped throughout the stimulus and delay intervals when mice used the integration strategy, but not when they resorted to the heuristic of relying on their bias (**Fig. 3g**). Indeed, during the heuristic strategy, choice-selective ALM activity showed a small increase at stimulus onset and remained mostly flat throughout — strikingly similar to what would be expected based on the psychophysical kernels observed for this strategy (see **Fig. 1i**). Second, the dynamics of choice-selective ALM activity observed for error trials were similar to those seen in the heuristic strategy. In contrast to correct trials, ramping was essentially absent during evidence processing, followed by a rapid surge at the end of the delay interval (**Fig. 3h**). Third, choice-selective ALM activity was stronger for decisions alternating the previous choice (**Fig. 3i**). Fourth, choice-selective ALM activity scaled with evidence strength (**Fig. 3j**). Finally, trial-by-trial variations in choice-selective ALM activity predicted RT (**Fig. 3k**). This latter aspect is a key signature of an accumulation-to-bound mechanism.^50^

In sum, ALM population activity contained two components: a global component that did not encode choice and a choice-selective component that reflected evidence integration. We next assessed the relationship between these two components of frontal cortical dynamics and pupil-linked, phasic arousal during decision-making, testing the hypothesis that the uncertainty-modulated arousal responses we observed during the task (**Fig. 2**) are driven by decision dynamics reflected in both components of ALM activity (**Fig. 3**).

### Both components of premotor cortical activity reflect pupil-linked, phasic arousal

An influential account holds that the cognitive recruitment of neuromodulatory systems – particularly the noradrenergic locus coeruleus – during decision formation is driven by top-down inputs from frontal cortical regions such as the anterior cingulate (ACC) or orbitofrontal cortex (OFC).^1^ ACC is closely linked to pupil dynamics^12,51,52^ and driven by cognitive conflict.^53^ Specifically, the ACC may respond to a conflict signal reflecting the total activity of neural accumulators supporting different options,^7^ which we approximated here in terms of the sum of left and right ALM activity (i.e., the global ALM component). Similarly, the models of the decision variable used for the computation of confidence and uncertainty in the static model of **Fig. 2** are commonly held to reflect the difference between competing accumulators, ^54–57^ which we approximated here in terms of the difference between ALM activity contra- and ipsilateral to the upcoming choice (i.e., the choice-selective component). Neurons in OFC encode either confidence or uncertainty signals that can be readily computed from this neural decision variable.^8,9^ These considerations predict a negative correlation between the choice-selective ALM component and pupil responses (i.e., the larger the ALM choice signal, the smaller uncertainty, in turn, exerting a smaller drive of arousal) and a positive correlation between the global ALM component and pupil responses. We next aimed to test these two predictions.

Trial-by-trial variations in both components of ALM dynamics were coupled with pupil responses during the task (**Fig. 4**), whereby the dynamics (and partially the sign) of this coupling differed markedly between the choice-selective and global components (compare **Fig. 4b** and **Fig. 4f**). Indeed, the coupling was positive for the global ALM component, indicating stronger activity predicting larger pupil responses during both, decision formation and after outcome delivery (**Fig. 4a-d**). This is consistent with the idea that a non-choice selective signal reflecting the overall accumulator activation (akin to a cognitive conflict signal) drives phasic arousal during and after decision formation.

**Figure 4.**
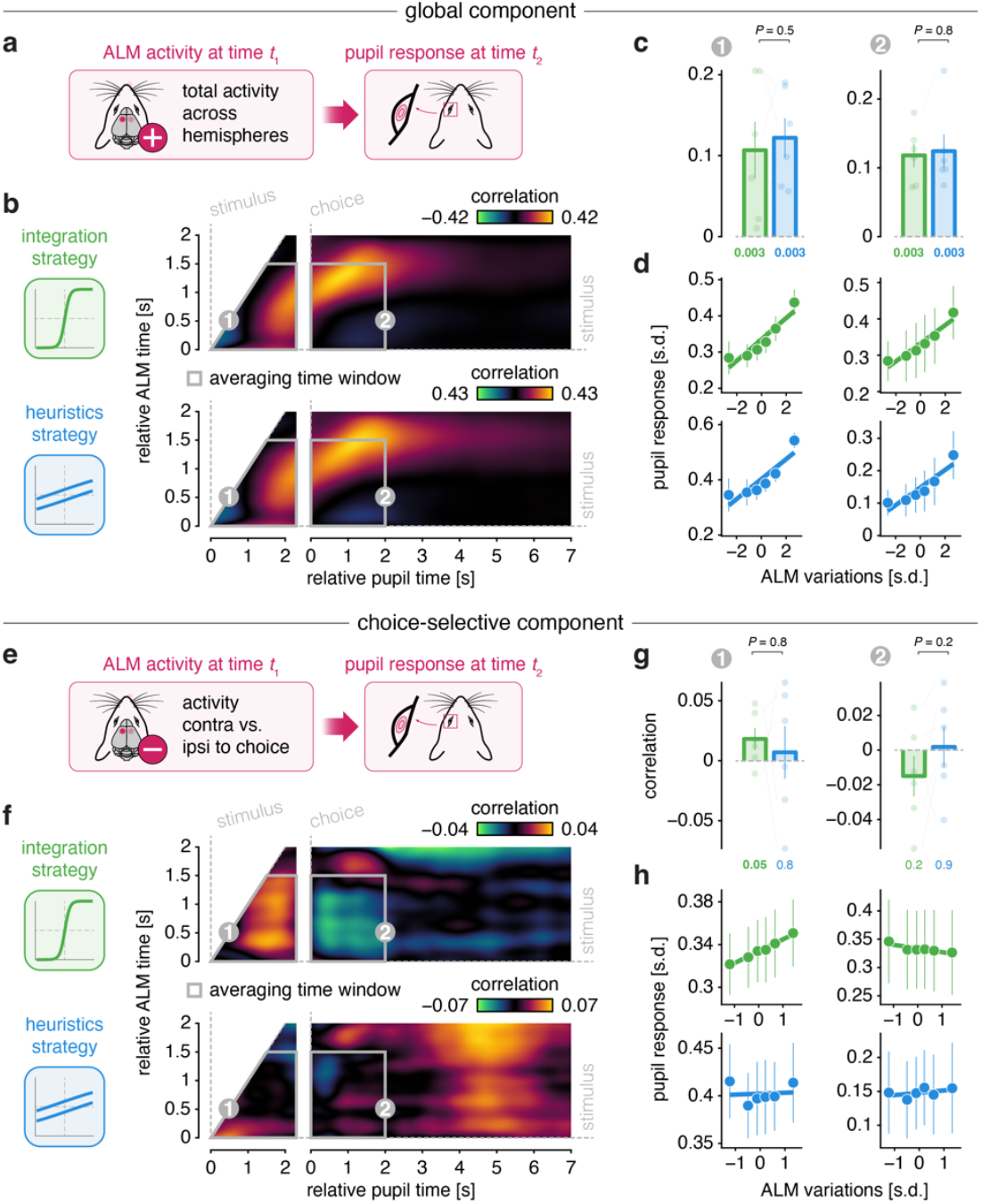
Pupil responses track moment-to-moment variations in two components of ALM activity. Trial-by-trial variations of ALM dynamics are related to pupil responses during the task. **a, e**, Illustration of analysis approach. Two components of ALM dynamics are studied: the choice-selective component (contra-minus ipsilateral to choice; a) and the global, non-selective component across hemispheres (left plus right; e). To account for the pupil’s slow dynamics, ALM and pupil are measured at different lags (denoted *t*_1_ and *t*_2_) relative to task events. **b, f**, Group mean correlation coefficient between ALM variations and pupil responses at all pairs of ALM and pupil timepoints (post ALM) separately for trials in which mice used and integration-based (top row) and heuristic-based strategy (bottom row). **c, g**, Group mean ± s.e.m. correlation coefficient between ALM variations and pupil responses in corresponding time-windows separately for the two different integration strategies. *P*-values are obtained from an exact permutation test. **d, h**, Group mean ± s.e.m. pupil response according to sextiles of ALM variations for the two different integration strategies.

The coupling between the choice-selective ALM component and pupil responses exhibited a more complex profile, with a positive correlation early during decision formation, followed by a negative correlation (blue blob) for the ALM signal during decision formation and the pupil response following choice report (**Fig. 4e-h**). While that later negative correlation is expected from a drive of arousal by an uncertainty signal reflecting the complement of the difference between cortical accumulators (see **Fig. 2a**), the positive correlation is not. In sum, pupil-linked, phasic arousal tracks the dynamics of decision-related signals in the frontal cortex, in a manner that is qualitatively in line with models of cognitive conflict.

### Pupil-linked, phasic arousal shapes subsequent choice behavior

Theoretical frameworks^1^ postulate that phasic arousal is not only driven by cortical decision dynamics, but that the resulting release of neuromodulators in the cortex in turn affects subsequent decision computation. We, therefore, reasoned that the pupil responses measured in one trial may predict alterations in choice behavior on the next trial. We tested this prediction focusing on two key aspects of choice behavior in our task: behavioral strategy (**Fig. 5a-d**) and serial choice bias (**Fig. 5e-h**).

**Figure 5.**
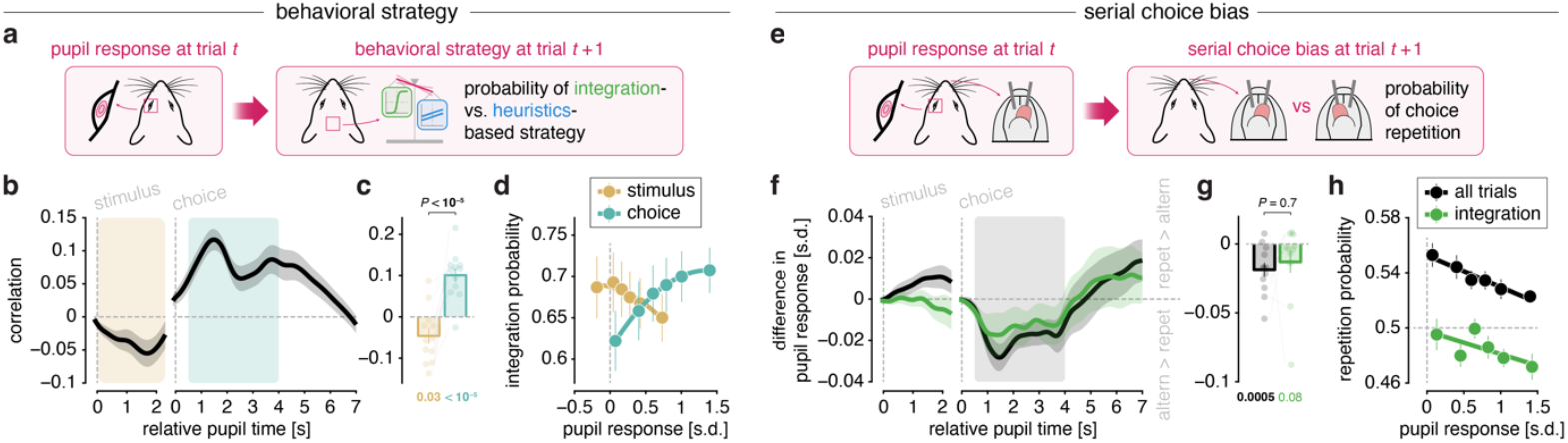
Pupil responses predict subsequent adjustment of choice behavior. **a**, Pupil response at trial *t* predicts behavioral strategy at trial *t* + 1 as estimated from the GLM-HMM (blue: heuristics, green: integration). **b**, Group mean ± s.e.m. time-course of the correlation coefficient between current pupil response to stimulus/choice and probability of using an integration strategy. **c**, Group mean ± s.e.m. correlation coefficient averaged in stimulus and choice time-windows (depicted as color shaded area in panel b). *P*-values are obtained from a random permutation test (10,000 permutations). **d**, Group mean ± s.e.m. probability of integration strategy (from the GLM-HMM) as a function of sextiles of pupil response to stimulus and choice. Lines are fits of a second-order polynomial. **e**, Pupil response at trial *t* is sorted according to the choice at trial *t* + 1. Choices at *t* + 1 are sorted according to whether they correspond to a repetition vs. alternation of choice at *t*. **f**, Group mean ± s.e.m. difference in current pupil response to stimulus/choice according to whether the next choice is a repetition (> 0) or an alternation (< 0) of the current choice. The difference is computed either from all trials (black) or only from pairs of trials in which the integration strategy was used (blue). **g**, Group mean ± s.e.m. difference in pupil response averaged in the choice time-window. **h**, Group mean ± s.e.m. repetition probability according to sextiles of pupil response to choice. Lines are linear fits.

Indeed, pupil responses predicted the strategy employed at the next trial as estimated by the GLM-HMM (**Fig. 1**), but with opposing signs during evidence processing and following choice (**Fig. 5b-d**). A larger pupil response following evidence onset favored the heuristic-based strategy while a larger response following choice report predicted the use of the integration strategy. Further, larger pupil responses following choice and reward predicted a reduction of choice repetition bias on the next trial (**Fig. 5f-h**), resembling similar observations in humans.^19^ This link to serial choice bias did not merely reflect the effect on behavioral strategy shown in **Fig. 5b-d**, which (as we have previously shown) went along with a switching in the propensity to repeat choices: there was a trend toward the same effect on trials where mice adopted the integration strategy (**Fig. 5g,h**; blue).

## DISCUSSION

We investigated whether and how pupil-linked arousal responses in mice reflect specific and rapid computations that unfold during perceptual decision formation. We trained mice in a challenging decision-making task which they solved by alternating between different strategies, driven either by heuristics (single-sample or side bias) or protracted evidence integration. The mice’s pupil responses during different periods of the trial scaled with decision uncertainty, mirroring patterns previously observed in humans. Moreover, moment-to-moment fluctuations in two distinct components of ALM population activity — a choice-predictive and a global component — were reflected in concurrent pupil responses. These arousal signals in turn predicted adjustments in behavioral strategy and the reduction of serial choice bias in the subsequent trial. Together, these findings establish a direct link between local cortical computations and the phasic recruitment of pupil-linked arousal systems, while demonstrating that this recruitment shapes subsequent adaptive behavior.

### ALM as a window into decision computations

Our analysis of ALM dynamics during decision formation extends prior work in two important aspects. Past studies have shown that the ALM exhibits preparatory motor activity that predicts the upcoming choice and reflects the outcome of sensory deliberation during delay periods.^43,44^ Here, we demonstrate that this preparatory signal ramps up during evidence accumulation and is graded by decision uncertainty. It exhibits stronger evidence-dependent ramping on correct trials and nearly absent ramping on error trials, which is a pattern consistent with the neural signature of a noisy accumulation process that sometimes reaches the wrong conclusion.

Second, we show that the cross-hemisphere imbalance of ALM population activity carries choice-predictive information that can be decoded from population recordings. This is analogous to mesoscopic decision signals that have been characterized non-invasively in humans.^27,58^ This observation establishes that choice-predictive cortical dynamics in mice can be detected without single-unit resolution, opening the door to wide-field imaging approaches that could be combined with circuit-level manipulations.

Third, we link decision-related ALM dynamics to the current behavioral strategy in mice. During the integration strategy, activity ramped progressively throughout stimulus and delay periods, which is consistent with ongoing accumulation of sensory evidence. In contrast, during the heuristic strategy ALM dynamics showed an initial transient followed by a plateau. This profile is strikingly consistent with a primacy-biased process, as if the animal’s decision was effectively made based on the first sample and then maintained rather than updated. This parallel between integration kernels and neural dynamics provides a rare instance of behavioral and neural signatures of strategy tracked together within the same framework.

When interpreting the ALM recordings, some technical aspects should be considered. Our dual-color recording strategy enabled us to spectrally separate the signals from the left and right ALM, thus avoiding crosstalk from the callosal projections of the other hemisphere. However, motion-related artifacts or hemodynamic responses might be a concern. On the one hand, we minimized the role of motion by focusing on periods preceding motor reports and using a signal reflecting the cross-hemisphere imbalance, hence canceling-out brain-wide signals such as motion artefacts. On the other hand, we minimized the contribution of hemodynamic effects by using bright indicators and confirming high signal-to-noise. Yet, we cannot entirely rule out that motion and hemodynamics might have contributed to ALM recordings.

### Task-related pupil responses report cortical decision signals

Prior work relating brain activity to pupil size has largely focused on slow, tonic variations in arousal, examining how baseline pupil diameter predicts engagement, vigilance, or neural gain in the subsequent epoch.^59–61^ These studies typically study how arousal state modulates cortical processing. Here, we invert this framing. By focusing on phasic, event-locked pupil responses and relating them to concurrent ALM dynamics within the same trial, we ask instead whether cortical processing leads pupil-linked arousal. Our results support this direction of influence: the magnitude of both ALM components — the choice-predictive imbalance and the total coordinated activity — predicted the magnitude of the pupil response to the same event. Interestingly, the overall activity of the anterior cingulate cortex, which is reciprocally connected to ALM and is involved in controlling and updating ALM, has also recently been shown to drive pupil-linked arousal.^52^ Our study extends this body of work by selectively investigating cognition-related cortical signals. While the correlational nature of our data does not permit causal conclusions, the specificity of these relationships, including their sensitivity to behavioral strategy and evidence strength, argues against a purely incidental association.

We further observed a rapid reconfiguration of arousal signaling across the trial. Stimulus-evoked pupil responses reflected an interaction between evidence and outcome consistent with pre-resolution decision uncertainty, while choice-evoked responses showed the opposite pattern. This sign-flip replicates a key feature of human pupil-linked arousa^l18,19^ and extends it to the mouse brain, suggesting that the computational logic governing phasic neuromodulatory recruitment is conserved across species. Notably, the modulation of the stimulus response by the evidence-by-outcome interaction is particularly diagnostic: it is the hallmark of a system that represents probabilistic belief states rather than simply responding to sensory intensity or motor effort. The fact that mice show this pattern strengthens the interpretation that their pupil fluctuations index genuine uncertainty-related computations rather than a simpler sensory or motor signal.

### Different timescales of pupil dynamics during cognitive tasks

Recent work in mice has focused primarily on tonic baseline pupil size as an index of behavioral state, demonstrating links to locomotion, arousal, and task engagement.^20^ By contrast, the phasic, event-locked component of pupil responses has received comparatively little attention in this species (see ^20,62,63^ nonetheless), in part because task designs have not always permitted the temporal isolation of stimulus- and choice-evoked responses. Our task design, with its explicit delay period separating sensory integration from motor report, was specifically constructed to allow this dissociation. The resulting data suggest that tonic and phasic pupil dynamics in mice carry complementary information: whereas tonic baseline may index the animal’s overall engagement or arousal level at trial onset, phasic responses reflect the specific computational events unfolding within the trial. The finding that phasic choice-evoked responses predict the behavioral strategy adopted on the subsequent trial, whereas baseline pupil at that subsequent trial likely reflects the residual dynamics of the prior response, illustrates how these two timescales of neuromodulatory signaling can interact in the shaping of sequential behavior.

### Broadcasting cortical computations across the forebrain

Choice-predictive activity in the form of contralateral motor preparation is not unique to ALM — similar ramping signals have also been identified in primary sensory cortices for instance.^32,41^ These signals are often interpreted as the consequence of top-down feedback from cortical regions involved in decision making and motor planning, reflecting the propagation of a forming decision backward through the cortical hierarchy. Our findings raise an alternative or complementary possibility: that phasic recruitment of brainstem neuromodulatory centers by active decision-making regions could itself contribute to the brain-wide broadcasting of preparatory signals, with ascending neuromodulatory projections modulating gain throughout the cortical hierarchy in concert with descending cortico-cortical signals. Testing this hypothesis directly will require combining our behavioral readouts with cell-type-specific recordings and manipulations of identified neuromodulatory populations.

## Conclusion

Our findings establish that phasic pupil-linked arousal responses in mice reflect moment-to-moment cortical decision computations and predict subsequent behavior. These findings bridge observations in humans with the mechanistic accessibility of the mouse, positioning the ALM-pupil relationship as a tractable system for dissecting how brainstem arousal circuits read out and broadcast cortical decision computations.

## ACKNOWLEDGEMENTS

We thank Carles Sindreu for supervising the design of the behavioral task. We thank Tiffany Oña-Jodar for sharing the design of the dual lick port and for advice on mouse training. We thank Cynthia Rais for help in managing the mouse colonies. We thank Stefan Schillemeit and Kathrin Sauter for extensive admin support, and Alexander Dieter for help in preparing the ethical protocol.

This work was supported by a *Humboldt Stiftung*, by the *Fondation Bettencourt Schueller* (both to M.M.) and by the *Deutsche Forschungsgemeinschaft (DFG, German Research Foundation)* SFB 936 - 178316478 - A7, Z3 (to T.H.D.) & B8 (to J.S.W.).

## AUTHOR CONTRIBUTIONS

Conceptualization: T.H.D., M.M., J.S.W.

Methodology: M.M., A.C., J.R., J.B.C., J.S.W., T.H.D.

Task design: A.C.

Software: M.M., J.B.C.

Validation: M.M.

Formal analysis: M.M.

Investigation: J.B.C., M.M.

Resources: A.C., J.R., J.S.W, T.H.D.

Data curation: J.B.C., M.M.

Writing — original draft: M.M.

Writing — review & editing: M.M., T.H.D., S.W., J.R., A.C., J.B.C.

Visualization: M.M.

Project administration: M.M.

Supervision: T.H.D., M.M., J.S.W.

Funding acquisition: J.S.W., T.H.D., M.M.

## METHODS

### Experimental model and subject details

#### Ethics

All experiments followed the 3R principles and were performed in compliance with German law according to the directive 2010/63/EU of the European Parliament on the protection of animals used for scientific purposes. The experiments performed in this study were approved by the Hamburg state authority for animal welfare under the license #005/2023. Sample size was determined according to past studies in the field.

#### Animals

Experiments were performed on 12 adult mice (4 females) aged 8 weeks at the start of the experiment (head-bar implantation). All mice were wild-types C57BL/6J obtained from *The Jackson Laboratory* (USA). Mice were kept in an animal room with controlled temperature (~22°C), humidity (~40%) and an inverted day-night cycle (12-12h), with free access to food. Following the surgical procedure, they were monitored daily until the end of the experiment. Mice recovered from the surgery for at least one week before they were habituated to the experimenter, the experimental setup, and to head fixation, in that order. After habituation, behavioral training commenced, followed by the data acquisition period.

### Surgical and molecular biology procedures

#### Surgical procedures

In the first batch of animals (*n* = 5 mice), the surgical procedure simply involved implanting a head-bar for head-fixation during the task. For the second animal batch (*n* = 7 mice), the surgical procedure also involved injection of viral vectors and fiber optic implantation above the injected areas to monitor brain activity during the task. Throughout the surgery, each animal was placed under general anesthesia through inhalation of isoflurane (induction: 2% for at least 5 minutes, maintenance: ~1.5%) mixed with air (RSA-A(H)-S; *RWD*, China). Analgesia was provided by subcutaneous injection of Buprenorphine (0.1mg × kg^−1^ in NaCl). Anesthesia and analgesia were confirmed by the absence of the hind limb withdrawal reflex, which was regularly checked throughout the surgery together with the breathing rate to fine tune isoflurane concentration. Body temperature (at 37°C) was maintained via a heating pad and monitored through a closed-loop system combining a rectal temperature probe and a heating controller (69027; *RWD*, China). In addition, temperature was also occasionally checked with an infrared thermometer (IR500-12S; *Voltcraft*, UK). After confirming anesthesia and analgesia, the animal was attached to a digital stereotactic surgical frame (100μm precision, 51725D; *Stoelting*, USA) using an auxiliary, blunted ear bar (EB-6; *Narishige*, Japan) to minimize damage to the animal’s ear tube and eardrum. The scalp was shaved using a trimmer (MT5531; *Grundig*, Germany) before being disinfected with iodide solution (Betaisodona; *Mundipharma*, Germany). Both eyes were then covered with eye ointment (Vidisic; *Bausch & Lomb*, Germany) to avoid drying. An antero-posterior (AP) skin incision was made (~1cm long), and the skull was cleaned from any remaining tissue. Anatomical landmarks bregma and lambda (taken as −4.1mm posterior to bregma) were aligned along the dorso-ventral (DV) and medio-lateral (ML) axes. Additional lateral landmarks (±2mm lateral to bregma) were further aligned along the DV axis. Two craniotomies were made (~500μm diameter) using a rotary micromotor (~6300rpm; K.1070; *Foredom*, USA) equipped with a steel drill (310 104 001 001 021; *Meisinger*, Germany), bilaterally above the ALM. For the injection of viral vectors, glass micropipettes were pulled from borosilicate glass capillaries (B100-20-10; *Sutter Instruments*, USA) using a micropipette puller (PC-100; *Narishige*, Japan). Each micropipette (one per injection site) was then filled with a suspension of adeno-associated viral (AAV) particles diluted in phosphate-buffered saline (PBS) solution before being slowly lowered into the ALM (±1.3mm ML, +2.3mm AP, −1.2/1.4/1.6/1.8mm DV to bregma). The viral suspension was slowly injected (at a rate of ~200nL × min^−1^) into the brain tissue using a custom-made manual air pressure system until the desired volume was reached (~100nL per site). The micropipette was kept in place for at least 1 min before it was slowly retracted. Once the two injections were completed, a 400μm diameter, 0.37NA mono fiber optic implant (MFC_400/430-0.37_1.2mm_ZF1.25(G)_FLT; *Doric Lenses*, Canada) was implanted vertically within each ALM (−1.5mm DV). Craniotomies were sealed and fiber optic implants were maintained in place using tissue adhesive (1469SB, Vetbond; *3M*, USA) which was also used to attach the skin to the skull. The skull and the base of the fiber optic implants were then covered with opaque dental cement (Super-Bond universal kit; *Sun Medical*, Japan). A head bar (200-200 500 2110; *Luigs & Neumann*, Germany) was finally implanted medially, posterior to the fiber optic implants when applicable, and further immersed in dental cement. An intraperitoneal injection of Carprofen (4mg × kg^−1^, diluted in NaCl) was performed as a post-surgery analgesic and anti-inflammatory treatment. The mouse was finally removed from the stereotaxic frame and allowed to recover alone in a dedicated cage partially placed on top of a heating pad. The animal was carefully monitored throughout the wakening stage and was placed back into its home cage after full recovery (as indicated by natural locomotion). Meloxicam (0.5mg × mL^−1^; Metacam; *Boehringer Ingelheim*, Germany) was mixed into softened food for the three days following surgery.

#### Genetic constructs and viral vectors

Two transgenes encoding genetically encoded calcium indicators (GECIs) were used: rAAV2/9-Syn-jGCaMP8m-WPRE (ITR-based titer = 5 × 10^12^ vg × mL^−1^; *AddGene* viral prep #162375; a gift from the GENIE project^65^) and rAAV2/9-hSyn-jRGECO1a-WPRE-SV40 (ITR-based titer = 2 × 10^14^ vg × mL^−1^; *AddGene* viral prep #100854; a gift from Douglas Kim & GENIE Project^66^).

#### Perfusion

After recording sessions were completed, the brains were collected for histological verification of the GECIs. Mice were first deeply anesthetized with intraperitoneal injections of ketamine and xylazine (respectively 180 and 24mg × kg^−1^ in NaCl). After the hind limb withdrawal reflex was fully abolished, the thorax was incised, the diaphragm opened and the thoracic cage removed. Mice were transcardially perfused, first with PBS (~100mL) to rinse out the blood, then with paraformaldehyde (PFA; 4%) in PBS (~100mL) to fix the tissue. The brain was then explanted and stored in PFA at 4°C in the dark for several weeks.

#### Immunohistochemistry

Brains were recovered and sliced coronally (70μm thickness) with a microtome (VT1000 S; *Leica*, Germany). The brain slices were incubated in normal goat serum (NGS; 10%) and Triton X-100 (0.3%) in PBS for 2h at room temperature to block unspecific antibody binding sites, followed by incubation for 2 days at 4°C in a NGS solution (10% NGS, 0.3% Triton X-100, in PBS) containing the primary antibodies. Afterwards, slices were rinsed 3 times for 5 minutes in PBS and subsequently incubated for 1 day at 4°C in a carrier solution (2% NGS, 0.3% Triton X-100, in PBS) containing the secondary polyclonal antibodies. jGCaMP8m was amplified using a chicken anti-GFP primary polyclonal antibody (1:1000 dilution; A10262; *Invitrogen*, USA) and a goat anti-chicken Alexa Fluor 488nm secondary polyclonal antibody (1:1000 dilution; A11039; *Invitrogen*, USA). jRGECO1a was amplified using a rabbit anti-DsRed polyclonal antibody (1:500 dilution; 632496; *TaKaRa*, Japan) and a goat anti-rabbit Alexa Fluor 546nm secondary polyclonal antibody (1:1000 dilution; A11035; *Invitrogen*, USA). After secondary antibody incubation, washing steps were repeated and brain slices were mounted on microscope slides using mounting medium (Fluoromount; *Serva*, Germany) and cover-slips (0.13-0.16mm thickness, 24 × 50mm; 1871; *Carl Roth*, Germany).

#### Microscopy

Full-view images of immunostained brain slices were obtained with an epifluorescence microscope (AxioObserver with Apotome 3; *Zeiss*, Germany) by stitching together individual tiles acquired with a 10× 0.45NA objective (Plan-Apochromat; *Zeiss*, Germany), controlled by the Zen software (v3.3; *Zeiss*, Germany). Presets corresponding to Alexa Fluor 488 and 555nm were chosen. Intensity of excitation light, exposure time and gain of the photomultiplier tubes were manually adjusted to each sample in order to maximize signal quality while avoiding image saturation. Image range, brightness and contrast were manually adjusted.

### Behavioral task

#### General description

Mice were trained to categorize the source (left or right) exhibiting the strongest auditory intensity by licking a reward port on the corresponding side. Correct responses were rewarded with a water drop (2.5μL; available for 2s) while incorrect responses were penalized by a time-out (2s). Lick spouts were only available at the day of the stimulus and delay periods (2s total). An inter-trial interval of 5s followed to ensure pupil could return to baseline before the next trial.

#### Generation of evidence levels

At each trial, a generative inter-aural level difference (ILD), expressed in decibel (dB), was drawn from a non-uniform trial distribution (favoring easy trials to maintain engagement). The ILD determines the generative difference in auditory intensity between the left and right speakers: the smaller the ILD, the lower the expected accuracy. The following ILD levels were tested: ±80dB, ±8dB, ±4dB, ±2dB and 0dB. By convention, negative/positive ILD indicate a larger intensity for the left/right speaker respectively. Trials with ±80dB ILD are unambiguous: only one of the two speakers is active. Trials with 0dB ILD are impossible to solve: both speakers are equally strong on average. In this latter case, reward was randomly allocated to one side with equal probability. At each trial, 10 samples were stochastically drawn from a bivariate gaussian distribution centered on the selected ILD level with a standard deviation of 1dB and within a ±4dB range. Each sample dictated the loudness of both speakers. Thus, the smaller the ILD, the higher was the likeliness that the sequence of 10 samples entailed inconsistent samples, whereby the loudest speaker was different from the globally loudest one.

#### Sound generation

Stimuli were made of 1s of white noise sampled at 44,100Hz and convolved with a 10-period sinusoid, giving rise perceptually to 10 bursts of noise. At each trial, this stimulus was then multiplied by the 10 generated samples of evidence to yield the final stimulus that was presented.

### Behavioral training

#### Training strategy

In line with standards in the field,^67–69^ mice were water deprived and water served as reward in the task. Each individual mouse was closely monitored for any sign of stress and body weight was monitored daily before each session. Daily water intake in the task was registered which, with weights, served to monitor the animal’s health and to fine-tune the maximum amount of daily intake, in order to maintain motivation while minimizing health impact. Weight loss was maintained above 80% of the initial weight, as measured on 5 consecutive days prior to the onset of water deprivation. Once habituation was complete, behavioral training started and consisted of the following four successive training phases.

#### Phase 1, lick teaching

During this phase, the animal first learned that water was dispensed in the rig. On the first day, the experimenter manually dispensed water with a 1mL syringe after the animal had explored the rig. On subsequent days, water was dispensed to either lick spouts while the animal was head-fixed. First, water was dispensed automatically on alternating sides. Then, water release was triggered by a lick. The animal was free to lick on either side but if it preferred one of the spouts more than the other, the preferred side was temporarily disabled to teach the animal that water is dispensed in an unbiased manner. This phase typically lasted 3 to 4 days.

#### Phase 2, categorization of unambiguous sounds

In this phase, the animal learned that the two lick spouts were associated with two different sound sources (left vs. right). Only unambiguous sounds were presented to the animal from only the left or the right speaker (±80dB ILD). A lick on the side of the active speaker was rewarded at each trial. In early trials, the mouse was allowed to correct itself if it initially licked the wrong side, so it could learn the correct association without experiencing strong penalties. As training progressed, incorrect choices were followed by a short timeout, and additional measures were used to reduce side bias (e.g., repeating the same side until the mouse adapted). An optional brief “no-lick” period before the sound was further introduced for impulsive animals. This phase continued until the animal reliably chose the correct side based on the sound with few misses, which typically occurred within 5 to 7 days.

#### Phase 3, waiting period

In this phase, the mouse continued to use the sound to decide whether to lick left or right, but now it had to wait before it could respond: the lick spouts were not available immediately, but were only presented following a delay period following sound onset. The delay period was gradually increased from no delay to the target value of 2 seconds. Thus, the animal had to learn to hold the sensory information in memory until the response opportunity appeared. As in the previous phase, training included basic anti-bias adjustments where necessary. This phase typically lasted for 10 to 15 days.

#### Phase 4, categorization of ambiguous sounds

In this phase, the task went beyond learning a simple left-versus-right sound-response. In addition, the animal started learning to base its choice on the amount of evidence in the stimulus. The goal was to train the mouse to integrate sensory information and choose the side that was better supported by all 10 samples. Trials began with an easy (±80dB ILD) 40-trial warm-up period to keep motivation high and bypass the impulsive behavior generally associated with session onset, followed by ambiguous (harder) stimuli as performance improved. The frequency of each of the 9 generative evidence strengths progressively changed as a function of the animal’s past performance to reach the final desired distribution of trial types. This phase continued until the animal performed reliably across all difficulty levels with minimal side bias, which was typically achieved within 5 days.

#### Data acquisition

When animals had reached full proficiency, the delay period was fixed at 1 second. After an initial 40 trials comprising only unambiguous sounds, trials were drawn from a non-uniform distribution over evidence strengths (easy trials more frequent). In each trial, the first lick was used to register the decision, and a 2s time-out followed in case of an incorrect choice. Behavioral data, pupil recordings, and brain activity were acquired in multiple sessions over a time period of several weeks with this version of the task without intervention of the experimenter during the sessions. A session was interrupted whenever the animal stopped licking in ~5 consecutive trials, or when the amount of desired water consumption was reached (tailored individually to each animal to maintain engagement in following days). On average, we collected 25 sessions per animal (range: 16-45) with an average of 398 trials per session (range across animals: 355-481).

### Experimental setup

#### Behavioral apparatus

The experimental setup was custom-made and consisted of a noise-reducing cabinet (9U; *Orion*, UK) with acoustic foam walls (MDM-40; *Monacor*, Germany) and stable, controlled ambient light (see pupil section below). The different hardware items listed below were placed on a breadboard (MB3030D/M; *ThorLabs*, USA) using head posts (*ThorLabs*, USA), in turn attached to a telescopic shelf (INT-TS-450; *Orion*, UK) to allow easy positioning of the animal and adjustments of the reward ports. Positioning was further facilitated by a camera providing a bottom view of reward ports and the animal’s mouth. Altogether, this ensured equal distance from the animal’s tongue to both reward spouts.

#### Synchronization

Task events were delivered using a Bpod finite state machine (r2.5, 1035; *Sanworks*, USA) and custom-made MATLAB code using the ‘TrialManager’ protocol, thereby ensuring minimal delay between trials. Continuous data streams (e.g., video) were recorded using Bonsai.^70^ Synchronization was ensured by soft codes sent from the Bpod to Bonsai at the onset of each trial through a local TCP protocol. Bpod, Bonsai and MATLAB were all operated by the same computer (4F856EA#ABD; *HP*, USA) running Windows 11 Education (10.0.22631 Build 22631; *Microsoft*, USA).

#### Lickometry

The water reward was delivered through custom-made, 3D-printed (with black PLA) lick ports. A lick was detected whenever the animal’s tongue broke into one of two photogates which were located in front of each lick port. Each photogate consisted of an infrared emitter (754-2220-ND; *DigiKey*, USA) and an infrared receiver (475-SFH310FA-2/3-ND; *DigiKey*, USA). Each pair of infrared dipoles was connected to a port interface board (1004; *Sanworks*, USA), itself under Bpod control.

#### Water delivery

Water delivery was triggered by the Bpod whenever the animal first licked on the correct side (determined by the generative ILD, or randomly for 0dB trials). Water drops were passed from a water tank (Perfusor 50mL; *B*|*Braun*, Germany) to the lick port through plastic tubing. The precise control of reward size was achieved by a solenoid valve (LHDA1231115H; *Lee Company*, USA) and regular calibration procedures.

#### Response window

The moment at which mice could report their decision was under experimental control. Specifically, reward ports were mounted on a custom-made, 3D-printed linear actuator (3170748; thingiverse.com), attached to a rotary servo motor (FS5106B; *FeeTech*, China). Reward ports were made available after a delay period of 1 second, which started after the sound presentation (1s) was over). This design facilitated measuring the pupil response specifically to the sound stimuli as well as the motor preparatory activity in absence of pronounced motor action. The servo motor of the water spouts was controlled by an analog input provided by a micro-controller board (Due; *Arduino*, Italy), which triggered changes in the analog signal in response to a digital input sent from Bpod in relevant trial periods.

#### Audio stimulation

Auditory stimuli were delivered by two ultrasonic speakers (L010; *Kemo Electronic*, Germany) which were placed at a 180° angle to the left and right of the animal at a distance of ~10cm to each side. The animal was head-fixed from the back to maintain free space between the speakers and the ears. The speakers were driven by an amplifier board (AMP2X15; *PUIaudio*, USA). For precise timing, sounds were pre-loaded on an SD card in trial dead times and triggered by a HiFi module HD (1033; *Sanworks*, USA), which was under control of the Bpod.

#### Audio calibration

Before training onset, each speaker was calibrated to 80dB using a microphone (UMIK-1; *miniDSP*, Hong Kong) logging air pressure levels (SPL; in dB) to the REW software (5.20.13; roomeqwizard.com) in response to a series of sounds whose intensity was varied along a grid of arbitrary units. SPLs were then fitted with an exponential function which was then used to estimate, for each speaker, the arbitrary level corresponding to a desired SPL level. Sound presentation, function estimation and inversion were all performed with custom-made MATLAB scripts.

#### Pupillometry

The animal’s left eye was illuminated with an infrared 850nm light source (IR30; *CMVision*, USA) and recorded with a monochrome camera (DMK 33UX249; *The Imaging Source*, Germany) equipped with an 8mm lens (TCL 0814 5MP; *The Imaging Source*, Germany) and a 695nm long-pass filter (FGL695M; *ThorLabs*, Germany). Baseline pupil size was maintained at an intermediate level (neither too constricted nor too dilated) by an ultraviolet LED (*Amazon*, USA) oriented towards the right eye and whose brightness level was controlled by a micro-controller board (Due; *Arduino*, Italy) through pulse width modulation. Video frames were acquired at a 30Hz frame rate and their corresponding time-stamps were recorded in Bonsai^70^ for further synchronization with other data streams and task events.

#### Calcium recordings

Activity of the left and right ALM was recorded with fiber photometry as bulk fluorescence emitted by two GECIs jGCaMP8m or jRGECO1a. Excitation and emission light travelled along the same optical path to and from the brain through the fiber optic implant. Excitation light was provided by two multi-color tunable LED light sources (pE-4000; *CoolLED*, USA) set to distinct excitation wavelengths (460-520nm, centered on 470nm for jGCaMP8m; 500-600nm, centered on 550nm for jRGECO1a – note that LED emission spectra are broad and were therefore additionally filtered by excitation filters). Excitation light power was set to 25-100μW on an animal-per-animal basis depending on overall fluorescence strength, and stimulation parameters were kept constant across acquisition sessions. Each light source was connected to a fluorescence mini-cube (FMC5; *Doric Lenses*, Canada) housing dichroic mirrors with built-in bandpass excitation (400-480nm for jGCaMP8m, 555-570nm for jRGECO1a) and emission (555-540nm for jGCaMP8m, 580-680nm for jRGECO1a) filters. The output light path of the fluorescence mini-cube was connected to the fiber optic implant by a low auto-fluorescence, 400μm diameter, 0.37NA patch-cord (MFP_400/440/1100-0.37_3m_FC-ZF1.25(F)_LAF; *Doric Lenses*, Canada) and a white 1.25mm zirconia mating sleeve (SLEEVE_ZR_1.25; *Doric Lenses*, Canada). Emitted fluorescence resulting from excitation of calcium indicators was detected with a photodetector system (gain = ×1; DFD_FOA_FC; *Doric Lenses*, Canada). Excitation lights and fluorescence read-outs were both controlled by a dedicated micro-controller board (pyBoard v1.1; *MicroPython*, USA) running a custom-modified version of the pyPhotometry software^71^ (with Python v.3.5) enabling control of two separate light sources and two photodetectors. To avoid the risk of cross-talk, we sequentially recorded from each of the two ALMs in a manner inspired from time-division multiplexing. Acquisition was done at a sampling rate of 90Hz for each of the two regions. In each acquisition sweep, background illumination was measured in a time window of 250μs immediately before excitation light was turned on for a duration of 750μs. Resulting fluorescence was measured in the last 250μs of the stimulation window and baseline-subtracted.

### Data analysis

#### General analysis approach

All analyses were made with custom-made scripts in MATLAB.

#### Hidden Markov model

The GLM-HMM was fitted using available *Python* code using a predefined number of 3 states.^28^ The GLM comprised two parameters: a coefficient for the effect of signed ILD on the probability of reporting a right choice and an intercept.

#### Psychophysical kernel

Kernels were obtained by averaging the ‘excess evidence’ (sample difference from mean) across trials, towards the chosen option separately for each choice and sample position and then (after sign-flipping) collapsing across choices.

#### Choice history kernel

Kernels were obtained by fitting a multiple logistic regression with 10 past choices and signed ILD (of current trial) as predictors.

#### Processing of lick recordings

Trials without motor report were removed from the analysis of lick trains. Licks correspond to periods during which the photogate was obstructed and are averaged across trials to yield a lick frequency metric, which corresponds to the proportion of trials during which the spout was licked at each particular time-point based on a 1000Hz sampling rate.

#### Processing of pupil recordings

Eight markers equally spaced along the pupil edge (left, left dorsal, dorsal, dorsal right, right, right ventral, ventral left) were used to define the pupil. Pupil markers were localized on each individual eye frame using DeepLabCut^72^. In short, a pre-trained 50-layer convolutional neural network (ResNet-50) was further trained for 1.000.000 iterations on 820 manually labeled eye frames from 26 sessions using a dedicated GPU (Quadro P2000 5GB GDDR5; *Nvidia*, USA). An ellipse was fitted on coordinates of pupil markers with confidence ≥ 90% using a least squares method. Pupil size was estimated as the diameter of the fitted ellipse, which corresponds to the average of the ellipse’s semi-minor and -major diameters. When less than 5 pupil markers were available, the corresponding pupil size sample was marked as missing. Moreover, samples with pupil size larger than 3 times the standard deviation of the pupil trace at that session, or with pupil derivative larger than 5 times the standard deviation of the pupil derivative trace at that session were labeled as bas. A 5-sample wide median filter was then applied to the resulting pupil size trace in order to remove local outlier samples. Epochs with more than 10% of missing samples were rejected (mean: 3.1% trials, 3.1-6.6% range) and other missing samples were linearly interpolated from neighboring, available samples. Finally, a 1.5Hz third-order Butterworth low-pass filter was applied to remove the remaining non-physiological variations. Baseline pupil size was estimated in a 500ms time-window immediately before event onset (stimulus or choice) and was regressed-out from the epoched pupil size to yield a phasic pupil response to task events. For averaging and statistical comparisons, pupil was averaged in a 0-2.3s time-window relative to stimulus and a 0.5-4s time-window relative to choice.

#### Preprocessing of calcium recordings

One mouse was excluded from the analysis calcium fluorescence due to weak fluorescence on one hemisphere, which was confirmed by cytotoxicity after histology. A 3Hz third-order Butterworth low-pass filter was applied on calcium traces to remove high-frequency noise. The traces were then epoched relative to stimulus onset. Photobleaching and reduction of dynamic range were accounted for by estimating the relative change in fluorescence as Δ*F*/*F*_0_ = (*F − F*_0_)/*F*_0_, where the baseline fluorescence (*F*_0_) was computed in a 500ms time-window immediately before stimulus onset. For averaging and statistical comparisons, calcium fluorescence was averaged in a 0-2s time-window relative to stimulus, corresponding to the both stimulus and delay periods prior to motor report.

#### Pupil-ALM correlation

The Pearson correlation coefficient between the ALM response to stimulus on one hand, and the pupil response to the stimulus or the choice. The correlation was computed for each pair *t*_ALM_ and *t*_pupil_ of twe two signals (with *t*_pupil_ ≥ *t*_ALM_). Correlation coefficient were then averaged in two-time windows ranging from 0-1.5s in ALM timing and 0-2s in pupil timing relative either to stimulus onset or choice.

#### Statistical testing

With the exception of single-mouse examples, all figures show results averaged across mice. Unless stated otherwise, all statistical tests were two-tailed, with a type-1 error risk set at α = 0.05. For all comparisons, a non-parametric, random permutation test was used: exact when based on calcium recordings (*n* = 6) and using 10,000 shuffles when used on behavior, licks and pupil data (*n* = 12).

